# Novel APC-like properties of human NK cells directly regulate T cell activation

**DOI:** 10.1101/016816

**Authors:** Jacob Hanna, Ofer Mandelboim

**Affiliations:** The Lautenberg Center for General and Tumor Immunology, Hebrew University–Hadassah Medical School, Jerusalem, Israel

**Author notes:** Present address: Department of Molecular Genetics, Weizmann Institute of Science, Rehovot 76100, Israel (Jacob Hanna). Address correspondence to: Ofer Mandelboim, The Lautenberg Center for General and Tumor Immunology, Hebrew University, Jerusalem 91120, Israel.

## Abstract

Initiation of the adaptive immune response is dependent on the priming of naive T cells by APCs. Proteomic analysis of unactivated and activated human NK cell membrane–enriched fractions demonstrated that activated NK cells can efficiently stimulate T cells, since they upregulate MHC class II molecules and multiple ligands for TCR costimulatory molecules. Furthermore, by manipulating antigen administration, we show that NK cells possess multiple independent unique pathways for antigen uptake. These results highlight NK cell–mediated cytotoxicity and specific ligand recognition by cell surface–activating receptors on NK cells as unique mechanisms for antigen capturing and presentation. In addition, we analyzed the T cell–activating potential of human NK cells derived from different clinical conditions, such as inflamed tonsils and noninfected and CMV–infected uterine decidual samples, and from transporter-associated processing antigen 2–deficient patients. This in vivo analysis revealed that proinflammatory, but not immune-suppressive, microenvironmental requirements can selectively dictate upregulation of T cell–activating molecules on NK cells. Taken together, these observations offer new and unexpected insights into the direct interactions between NK and T cells and suggest novel APC-like activating functions for human NK cells.

**Nonstandard abbreviations used:**

ANK: activated NK
dNK: decidual NK
HA: hemagglutinin
ImDC: immature DC
MDC: mature DC
MFI: mean fluorescence intensity
NCR: natural cytotoxicity receptor
TAP2: transporter-associated processing antigen 2
UaNK: unactivated NK

## Introduction

The immune system displays efficient protective capabilities against an impressive array of pathogens and tumor cells (1). The invading pathogens are either kept in check or eliminated first via cells of the innate immune system and then by cells of the adaptive immune system (2). The initial action of the innate immune system will also shape the nature of the adaptive immune response that follows (3). Cytokines are secreted, and APCs capture and process the invading pathogen via multiple mechanisms and activate T lymphocytes (4, 5). The naive and memory T cell responses to antigen are directed by immunogenic stimuli that can be divided into 2 categories: first, TCR ligation mediated by peptide-MHC complexes; and second, costimulation signal conferred by counter-receptor expressed on APCs, which is mandatory for efficient priming and avoidance of T cell anergy (2).

NK cells that are part of the innate immune system are among the first immune effector cells to arrive at inflammation sites (6). In addition, NK cells have extensive trafficking capabilities and are found in lymphoid and nonlymphoid organs (7, 8). The major function of NK cells is to kill their target cells. NK cells also indirectly shape the nature of the adaptive immune response, mainly by producing profound amounts of cytokines that result in activation of monocytes and priming of DCs (3, 9). On the other hand, it was also suggested that NK cells can attenuate adaptive immune response by killing autologous immature DCs (10). Still, however, unlike for other key innate immune cells that are recruited early to inflammation sites, a direct role for NK cells as APCs or TCR costimulators remains to be characterized.

In this study we characterized the expression and function of MHC class II proteins and ligands for TCR costimulatory molecules on human NK cells following activation. More importantly, we extend the immunological relevance of these observations by further estimating NK capabilities and novel mechanisms to capture, process, and present antigens. Potential implications of our findings for fundamental immunological scenarios are discussed.

## Results

### High-throughput proteomic analysis of NK cell membrane–enriched fractions

The question of whether human NK cells possess APC-like properties was raised after experiments involving high-throughput proteomics techniques to obtain information about proteins in human NK cells. We prepared membrane enriched fractions from purified unactivated NK (UaNK) and activated NK (ANK) cultures. The complex membrane protein mixtures were separated into 20 molecular mass fractions by 1-dimensional gel electrophoresis followed by excision of equally spaced bands (Figure 1). The spectra were searched against a human database using probability-based scoring. As expected, the visually distinct alteration in the pattern of ANK membrane proteins (Figure 1) yielded a different identified protein repertoire for each of the fractions. This query resulted in a list of over 2,000 proteins identified from ANK and UaNK cells (data not shown). Multiple NK receptors were present in both ANK and UaNK cell membrane–enriched fractions (Table 1) and correlated with the specific phenotype of the NK cells analyzed (Figure 2A). Several interesting proteins identified from ANK cell membrane–enriched fractions are key players in stimulating T cells. ANK cells expressed significant levels of MHC class II and ligands for TCR costimulatory molecules (B7-H3 and CD70) (Table 1), while data analysis of UaNK cells yielded lower levels of MHC class II–derived peptides and no detection of either CD70 or B7-H3 molecules (Table 1). We verified the proteomics data by flow cytometry. MHC class II was expressed at low levels on UaNK cells (Figure 2B), confirming previous reports (11), while ANK cells expressed significant levels of CD70 and MHC class II (Figure 2B). B7-H3 expression was validated on NK cells via RT-PCR analysis that yielded specific detection only in ANK cells (Figure 2C). In addition, ANK cells significantly expressed OX40 ligand, CD80, and CD86 (Figure 2B).

**Figure 1.**
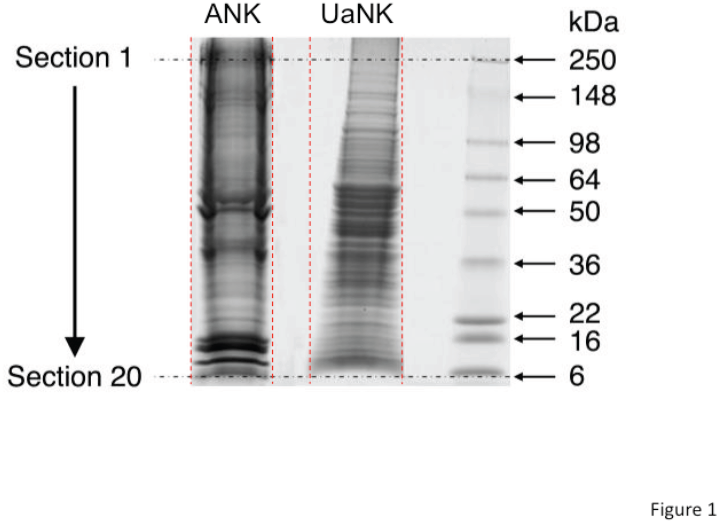
One-dimensional gel separation of NK membrane protein fractions. One hundred micrograms of UaNK and ANK cell membrane-enriched fractions were separated on 4-20% Tris-Glycine gel. Twenty equally spaced sections between 250 and 6 kDa were excised from each fraction and were subsequently used for proteomic analysis. Red dotted vertical lines indicate cropping borders for each lane.

**Table 1.**
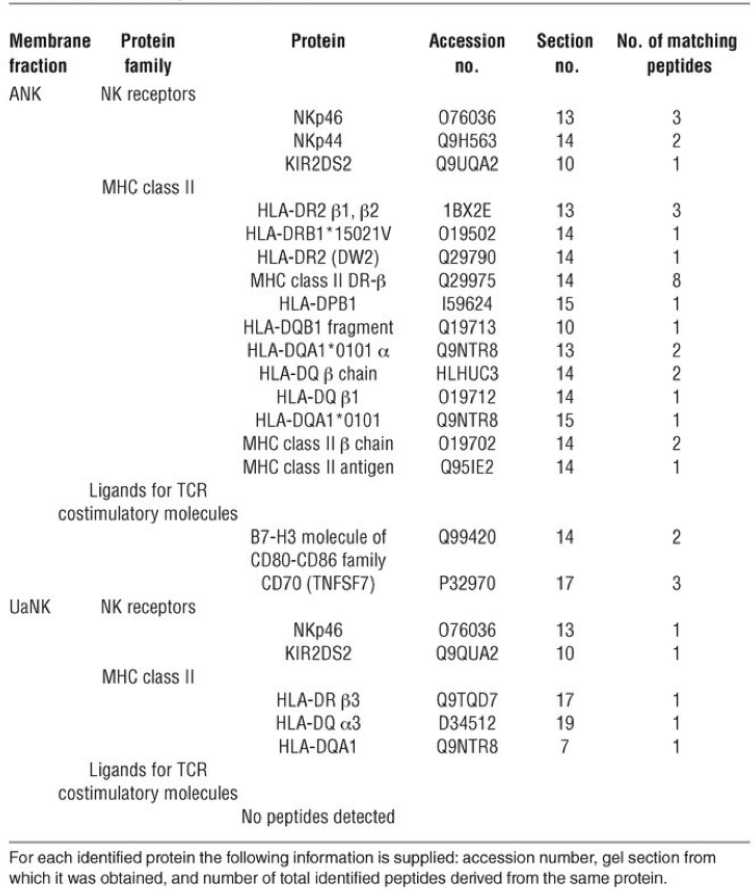
Identified NK receptors, MHC class II proteins, and ligands for TCR costimulatory molecules, based on mass spectrometry/mass spectrometry proteomic analysis of ANK and UaNK cell membrane-enriched protein fractions

**Figure 2.**
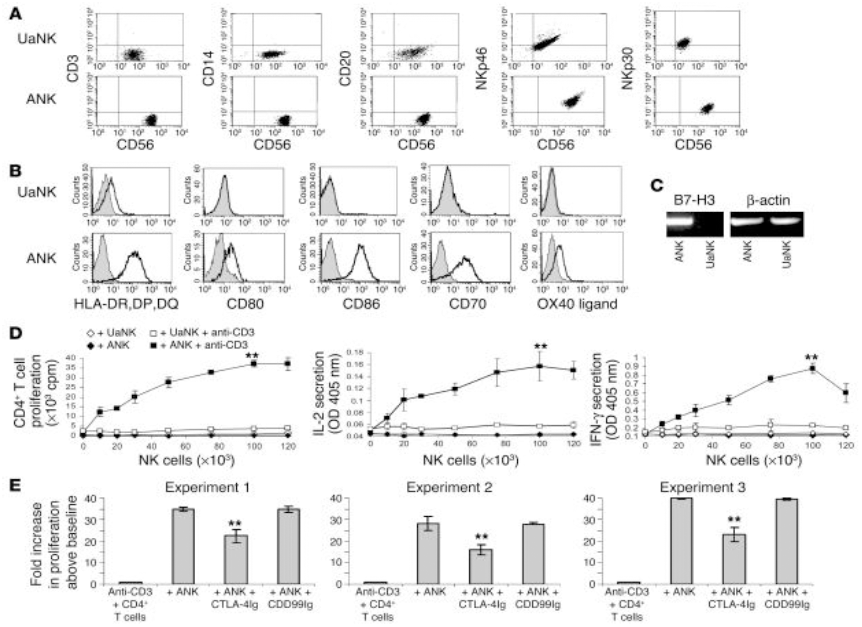
Assessment of the functionality of NK cells as costimulators for T cells. **(A)** Representative characterization of UaNK and ANK cells used in this study. Cells were stained for CD56 and for CD3, CD14, CD20, NKp46, or NKp30. (B) UaNK and ANK cells were stained for the expression of MHC class II or ligands for TCR costimulatory molecules. (C) RT-PCR of B7-H3 expression on ANK and UaNK cells. (D) Proliferation of CD4^+^ T cells in the presence of irradiated ANK or UaNK cells with or without suboptimal levels of anti-CD3. Proliferation was determined after 48 hours. Prior to cell harvesting, supernatant from each well was taken for measurement of IL-2 and IFN-γ. (E) Data obtained from 3 independent experiments are presented to show the effect of CTLA-4Ig fusion protein or CD99Ig fusion protein (25 μg/ml). Values are mean ± SD for triplicate samples. **P < 0.01 by Student’s t test. One representative data set is shown out of 3-12 obtained.

To rule out the possibility that our polyclonal culturing conditions for ANK cells were selectively enriching a minor subset of NK cells with APC-like properties, we characterized the expression of all molecules on activated NK clones. Our analysis showed that, irrespective of the inhibitory-receptor repertoire of each clone, all of the NK clones studied expressed similar levels of MHC class II and ligands for TCR costimulatory molecules (except for CD70, which was expressed on about 65% of the clones) (Table 2).

**Table 2.**
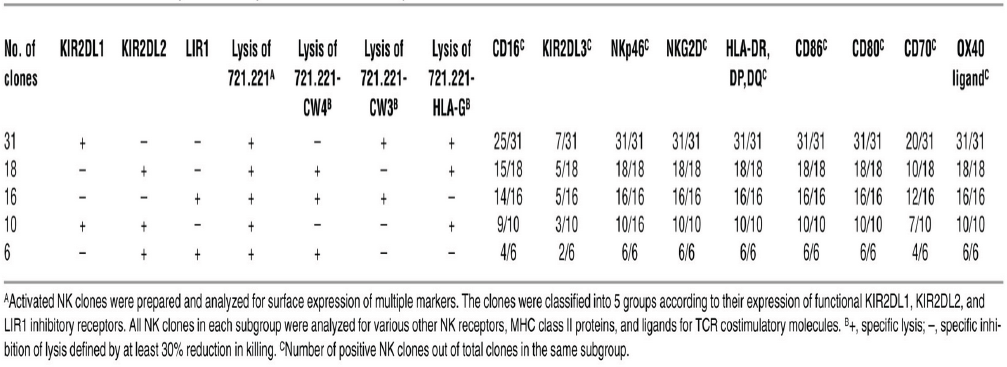
Characterization of MHC class II proteins and ligands for TCR costimulatory molecules on activated NK clones.

### ANK cells costimulate TCR responses

The functional importance of these costimulatory molecules on NK cells was examined using low concentrations of plate-bound anti-CD3 mAb and subsequent addition of irradiated UaNK or ANK cells. Remarkably, incubation with ANK cells resulted in a dose-dependent enhancement of CD4^+^ T cell proliferation and IL-2 and IFN-γ secretion (Figure 2D) only when CD3 cross-linking was applied. The maximal effect was observed at an NK/T cell ratio of 2:1, while UaNK cells did not significantly enhance CD4^+^ T cell activation beyond the basal levels (Figure 2D). Preincubating ANK cells with CTLA-4Ig protein that blocks CD80 and CD86 interactions resulted in a reduction of approximately 35% of the ANK-mediated enhancement of T cell proliferation (Figure 2E). The control CD99Ig fusion protein had no effect. A reduction by approximately 55% in cytokine secretion following CTLA-4Ig treatment was also obtained (data not shown). Similar results were obtained when CD8^+^ T cells were studied (data not shown). Collectively, these findings provide a proof of concept for the ability of ANK cells to function as accessory cells that can costimulate T cells following initiation of TCR activation.

### Characterization of MHC class II–restricted interactions between human ANK and CD4^+^ T cells

The significant expression of MHC class II and costimulatory molecules provided us with justification to substantiate the possible role of ANK cells as APCs in a specific MHC class II–restricted system. We generated a CD4^+^ T cell clone (OFR3 clone), from an HLA-DR4^+^ donor, that is specific for the influenza hemagglutinin-derived (HA-derived) peptide HA_306–318_ presented on HLA-DR4 (12, 13). The specificity of this clone was confirmed by binding to HLA-DR0401 tetramers loaded with HA peptide, but not to those loaded with control ospA peptide (Figure 3A). Antigen-specific proliferation of the T cell clone was observed when UaNK and ANK cells derived from the same donor were pulsed with the HA_306–318_ peptide (Figure 3B). The lower proliferation levels obtained with the UaNK cells can be attributed to lower levels of MHC class II and costimulatory molecules found on these cells (Figure 2B). Similar results were achieved when APC abilities were explored, as ANK cells were able to process recombinant HA protein and induce antigen-specific proliferation of OFR3 T cells (Figure 3C). This antigen-specific proliferation by ANK cells was partially dependent on CD80 and CD86 expression, since a significant reduction in proliferation was observed following CTLA-4Ig treatment (Figure 3, B, and C). We next compared the in vitro APC properties of NK cells to those of immature and mature DCs (ImDCs and MDCs, respectively) in our restricted experimental model. As expected, ANK cells were less potent in inducing OFR3 T cell proliferation following pulsing with either HA peptide (Figure 3B) or recombinant HA protein (Figure 3C), but still, OFR3 stimulation by ANK cells was highly significant. We ruled out the possibility that these results were due to contaminating B lymphocytes or monocytes, as the ANK lines used were consistently 100% pure (median purity = 100% ± 0% in 18 different cultures examined) and fresh NK preparations were uniformly greater than 98% pure (median purity = 98.7% ± 0.6% in 13 different cultures examined) and contained no detectable CD14^+^ or CD20^+^ cells (representative staining in Figure 2A).

**Figure 3.**
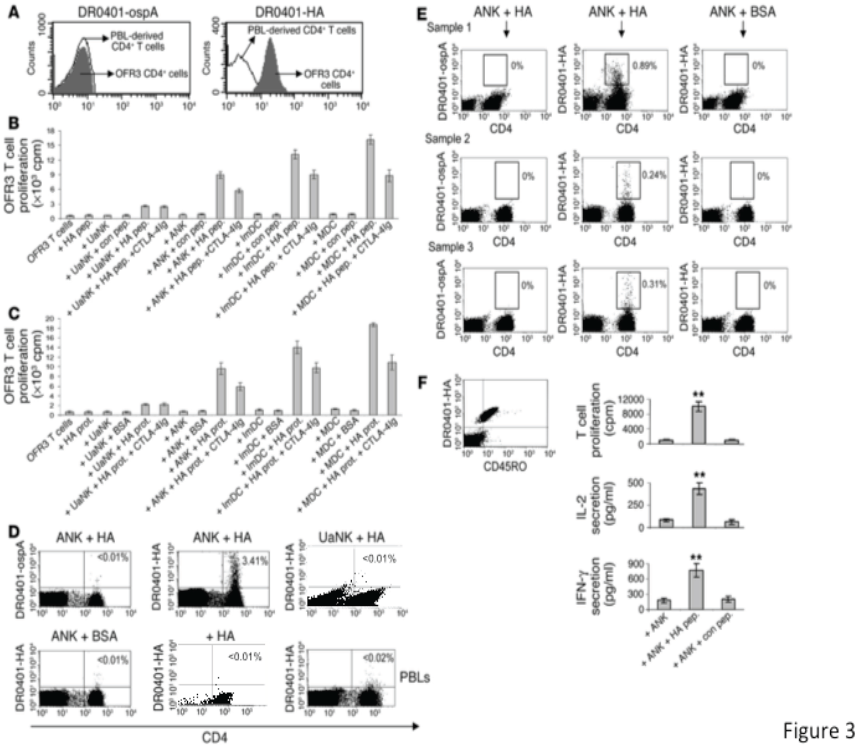
MHC class II-restricted interactions between ANK and CD4^+^ T cells. **(A)** OFR3 CD4^+^ T cells were stained with PE-labeled DR0401 tetramer loaded with HA (DR0401-HA) or control peptide (DR0401-ospA) (gray histograms). Staining of freshly isolated CD4^+^ T cells with the same tetramers was used as a background setting (white histograms). (B and C) All APC_T cell incubations applied in the following experiments were maintained for 48 hours followed by addition of thymidine for an additional 24 hours. (B) Irradiated cells (50 × 10^3^) were incubated with either HA_306–318_ (HA pep.) or control peptide (con pep.) (150 ng/ml) and added to the OFR3 T cell clone (50 × 10^3^ cells per well). (C) Antigen-processing capability was tested by measurement of OFR3 clone proliferation in the presence of 150 ng/ml recombinant HA or BSA proteins (prot.). The contribution of CD80 and CD86 to OFR3 T cell activation was evaluated by pretreatment of APCs with 25 ng/ml CTLA-4Ig. (D) CD4^+^ T cells isolated from HLA-DR4^+^ donors were incubated for 10 days with UaNK or ANK cells pulsed with HA protein or BSA (indicated on the top of each panel), or only with recombinant HA protein. Cells were stained for CD4 and DR0401-HA or control DR0401-ospA tetramers. The percentage of tetramer-positive fraction out of total CD4^+^ T cells is indicated. Staining of PBLs from the same donor is shown as a control. One representative staining obtained from a single donor out of 3 donors examined is shown. (E) Naive umbilical cord-derived CD4^+^ T cells were cultured with HA- or BSA-pulsed ANK cells for 14 days and stained with DR4 peptide-loaded tetramers. (F) DR0401-HA_positive umbilical cord-derived CD4^+^ T cells were characterized for CD45RO and antigen-specific effector responses. At day 25 of coculturing, 150 × 10^3^ cells from each donor were incubated for 96 hours with 150 × 10^3^ irradiated autologous ANK cells either unpulsed or pulsed with HA or control peptide, in the presence of 2 μCi of [^3^H]thymidine per well. Prior to harvesting, supernatant was taken for measurement of IL-2 and IFN-γ. One representative characterization from a single donor out of 3 is shown. Values are mean ± SD for triplicate samples. **P < 0.01 by Student’s t test.

We next studied the ability of ANK cells to expand PBL-derived memory CD4^+^ T cells specific for HA. We isolated CD4^+^ T cells from 3 HLA-DR4^+^ donors previously exposed to influenza virus. These cells were cultured with autologous ANK cells preincubated with HA protein or BSA and were subsequently stained for the detection of CD4^+^ T cells that specifically bound DR0401–HA peptide tetramers. Indeed, after culturing with ANK cells pulsed with HA protein, the frequency of DR0401-HA tetramer–positive CD4^+^ T cells, but not of control DR0401-ospA tetramer–positive cells, was specifically increased compared with the frequency after culturing with BSA-pulsed ANK cells, HA-pulsed UaNK cells, or freshly isolated CD4^+^ T cells (Figure 3D). To rule out the possibility that other contaminating APCs were present, we incubated purified CD4^+^ T cells with HA protein in the same culturing conditions and observed no expansion of HA-specific CD4^+^ T cells (Figure 3D). The observation that UaNK cells pulsed with HA protein failed to expand the antigen-specific T cells might be attributable to their reduced MHC class II expression and lack of T cell–costimulating molecules as shown earlier (Figure 2B).

More importantly, we were interested in evaluating the ability of human ANK cells to prime HA-specific naive CD4^+^ T cells. Since the fetal immune system has never coped with influenza infection before, we used as a model umbilical cord–derived naive CD4^+^ T cells obtained from HLA-DR4^+^ healthy neonates. Remarkably, in 3 different donors, DR0401-HA tetramer–positive CD4^+^ T cells could be detected following incubation of these naive cells with HA-pulsed autologous ANK cells, but not following incubation with BSA-pulsed autologous ANK cells (Figure 3E). The observed staining was specific, since no detection was observed with DR0401-ospA control tetramer. We further expanded these cultures and checked whether the primed T cells were functional. Remarkably, in all umbilical cord–derived CD4^+^ T cell cultures, DR0401-HA–positive cells were CD45RO^+^, and we were able to evoke antigen-specific proliferation and cytokine secretion when HA peptide–pulsed ANK cells, but not when unpulsed ANK cells or ANK cells pulsed with control peptide, were reintroduced to these cultures (Figure 3F). We could not evaluate UaNK cells’ ability to prime naive T cells, because of the limited amount of cells that could be obtained from human umbilical cord samples. However, since these cells were not able to expand memory HA-specific CD4^+^ T cells in cultures derived from adult donors (Figure 3D), it is most likely that UaNK cells are not able to prime naive CD4^+^ T cells.

NK cells acquire functional APC-like properties after target-cell killing. Can NK cells acquire APC-like properties after activating receptor–mediated recognition and killing of their target cells? Freshly isolated NK cells were incubated for 72 hours with various target cells known to be efficiently lysed by them. Strikingly, MHC class II and CD86 were specifically induced on NK cells following killing of multiple susceptible target cells (Figure 4A). By contrast, no upregulation was observed following incubation with NK lysis–resistant P815 cells; this implicates target cell killing as an additional important pathway for acquiring of TCR costimulatory molecules and upregulation of MHC class II.

**Figure 4.**
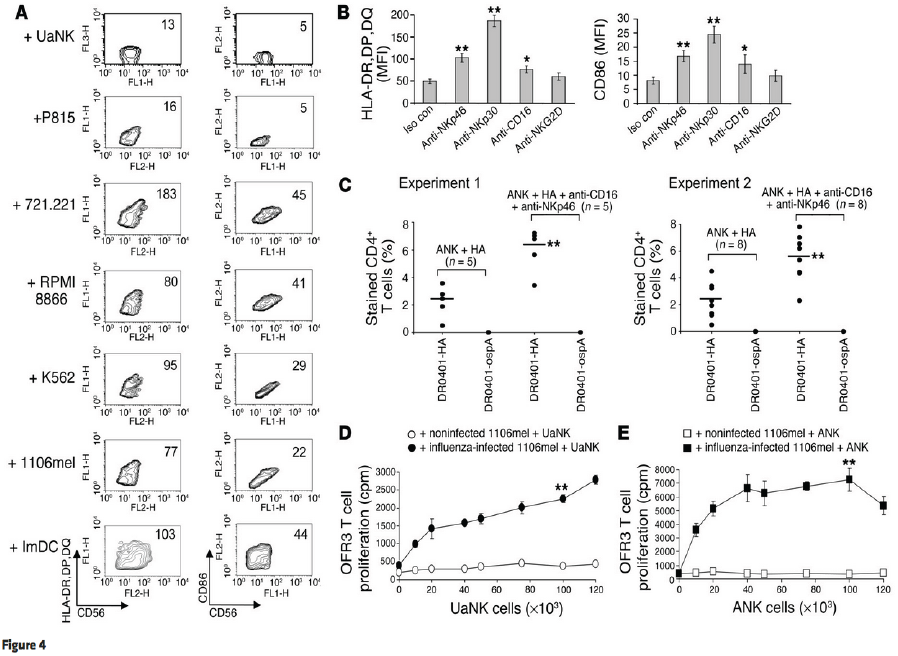
Enhanced APC-like properties of NK cells after lysis receptor-mediated activation. **(A)** UaNK cells (100 × 10^3^) were cocultured with multiple target cells at an effector/target ratio of 1:1 for 72 hours. NK cells were purified using magnetic beads and stained for CD56 and either HLA-DR,DP,DQ or CD86. MFI for the latter 2 markers is indicated in each panel. (B) UaNK cells were cultured in triplicate for 72 hours in RPMI medium plus 10% human serum and 30 U/ml human IL-2 and in the presence of the indicated soluble mAbs and were stained for expression of HLA-DR,DP,DQ and CD86. Data are shown as MFI ± SD. Iso con, isotype-matched control. (C) Circulating CD4^+^ T cells (4 × 10^6^) isolated from HLA-DR4^+^ donors were incubated with 1 × 10^6^ ANK cells pulsed with 150 ng/ml HA protein per well in 24-well plates. On day 4, half of the wells in each experiment were supplemented with 5 μg/ml of anti-CD16 and anti-NKp46 mAbs. On day 10, cells from each well were stained for CD4 and DR0401-HA or control DR0401-ospA tetramers. The percentage of tetramer-positive fraction out of total CD4^+^ T cells stained is indicated. (D and E) UaNK cells (D) or ANK cells (E) (0.5 × 10^6^ each) were incubated with noninfected or influenza virus-infected 1106mel cells at an NK/target ratio of 10:1 for 72 hours. The NK cells were repurified with magnetic beads, irradiated, washed, incubated with OFR3 clone cells for 48 hours, and pulsed with thymidine for the last 24 hours. *P < 0.05; **P < 0.01 by Student’s t test. Data are representative of 2-4 independent experiments.

To further support the role of specific NK-mediated cytotoxicity in regulation of MHC class II and CD86, we incubated freshly isolated NK cells with antibodies against NKp30, NKp46, CD16, or NKG2D (Figure 4B). Cross-linking of all receptors, except for NKG2D, led to upregulation of MHC class II and CD86. Our finding that anti-NKG2D mAb is not sufficient to induce expression of these molecules might be attributed to previously observed assay limitations (14).

Activating receptors on DCs, such as toll-like receptors, have been shown to enhance antigen-specific responses in vitro via cytokine production (15). As NKp46 and CD16 cross-linking on NK cells is known to regulate cytokine secretion, we wondered whether a similar role might be played by these receptors in influencing T cell responses. We incubated CD4^+^ T cells with HA protein–pulsed ANK cells, and on the fourth day, half of the wells were supplemented with a combination of anti-CD16 and anti-NKp46 cross-linking mAbs. Interestingly, such treatment enhanced the expansion of DR0401- HA tetramer–positive CD4^+^ T cells more than 2-fold, as detected at day 10 of coculturing (Figure 4C).

Next, we explored whether NK lysis of influenza-infected cells can result in the acquisition of HA by NK cells and subsequently induce specific T cell proliferation. Remarkably, both UaNK (Figure 4D) and, more potently, ANK cells (Figure 4E) that killed influenza-infected targets were able to specifically activate the HA-restricted T cell clone. Addition of influenza-infected target cells alone (data not shown) or NK cells that were incubated with noninfected target cells failed to demonstrate a similar effect (Figure 4, D, and E). These results suggest that NK cells can first kill the invading pathogen, then become activated, internalize, process pathogen-derived antigens, and stimulate CD4^+^ T cell responses.

### Activating NK receptors can internalize into MHC class II peptide loading compartments

It was recently shown than antigen transfer to NK cells occurs following target-cell recognition (16). This raises the question of whether NK activating receptors are involved in antigen capturing following such specific interactions. We observed a time-dependent internalization of NKp30, NKp46, NKG2D, and CD16 following antibody cross-linking and subsequent incubation at 37°C (Figure 5A). To show that ligation of NK activating receptors with their natural ligands can be sufficient to cause downregulation, we incubated ANK cells with 721.221 cells that express unknown cellular ligands for NKp30 and NKp46, but not for NKG2D (17). Indeed, a rapid downregulation of NKp30 and NKp46 receptors was observed following incubation with 721.221 cells, while NKG2D levels remained constant (Figure 5B). The NKG2D downregulation was studied by incubation of ANK cells with C1R cells transfected with NKG2D ligands MIC1A, MIC1B, and ULBP1 (18). A specific downregulation of NKG2D was observed following interaction with all of these ligands (Figure 5C). To determine whether this downregulation in NK activating receptors after interaction with their ligands resulted from receptor internalization, NK cells were fixed, permeabilized, and stained for the presence of intracytoplasmic NK activating receptors. No significant alteration was observed in the total amount (surface and intracellular) of NKp30, NKp46, and NKG2D detected by staining of permeabilized NK cells (Figure 5, D and E). Taken together, these results show that specific ligand-mediated modulation of NKp46, NKp30, and NKG2D surface expression is most likely caused by internalization.

**Figure 5.**
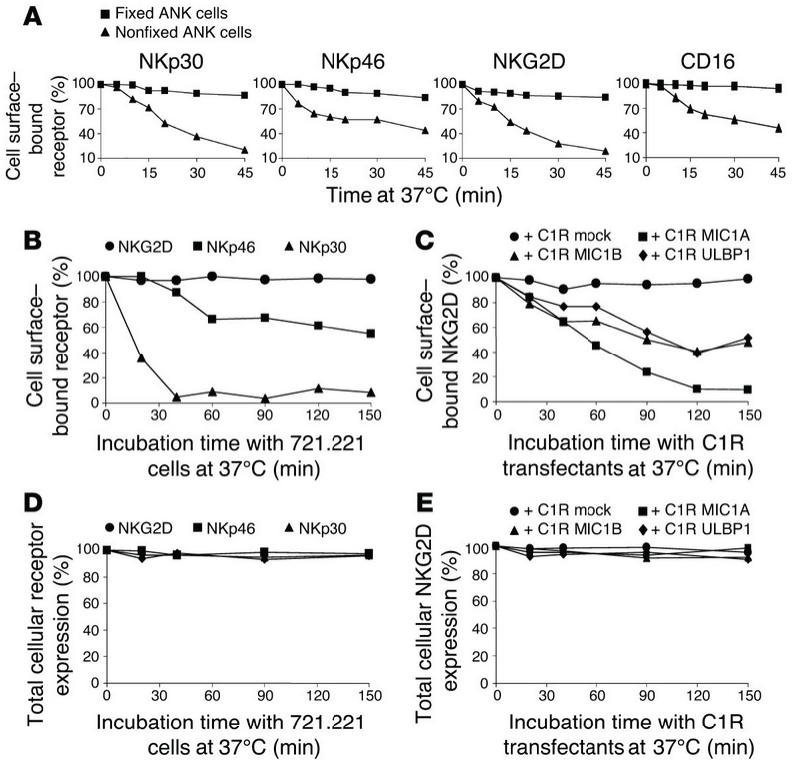
Characterization of activating NK receptors’ internalization following target recognition. **(A)** Time-dependent percentages of surface levels of NKp30, NKp46, NKG2D, and CD16 receptors on fixed and nonfixed ANK cells. (B) Modulating NKp30 and NKp46 surface expression following ligand-specific recognition. ANK cells (50 × 10^3^) were incubated with an equal number of 721.221 cells at 37_C for 150 minutes. The figure shows time-dependent changes in the percentages of surface levels of NKG2D, NKp46, and NKp30. (C) Down regulation of NKG2D following interaction with its ligands. ANK cells (50 × 10^3^) were incubated with an equal number of C1R transfectants, or with control C1R (C1R mock). The figure shows time-dependent changes in the percentages of expression of NKG2D following specific interaction with its ligands. (D and E) The same coculturing conditions described in B and C, respectively, were applied. Subsequently, NK cells were permeabilized and stained for total expression (intracellular plus surface) of NKp30, NKp46, and NKG2D. The figure displays total cellular expression levels of these receptors following ligation with their specific ligands.

The next step was to elucidate whether NK activating receptors intersect, upon internalization, with endosomal-lysosomal compartments, where conventional peptide loading of MHC class II molecules occurs (4, 5). We labeled ANK cells with a marker that stains acidified endosomal-lysosomal compartments, then subsequently incubated ANK cells with antibodies against CD16, which is known to internalize into such MHC class II peptide loading compartments on DCs (Figure 6A) (4), and with NKp46, NKp30, or NKG2D, and induced their internalization following cross-linking. Confocal microscopy experiments showed surface expression of these molecules prior to incubation at 37°C, whereas after 30 minutes of incubation all 4 receptors internalized into endosomal-lysosomal compartments (Figure 6A).

**Figure 6.**
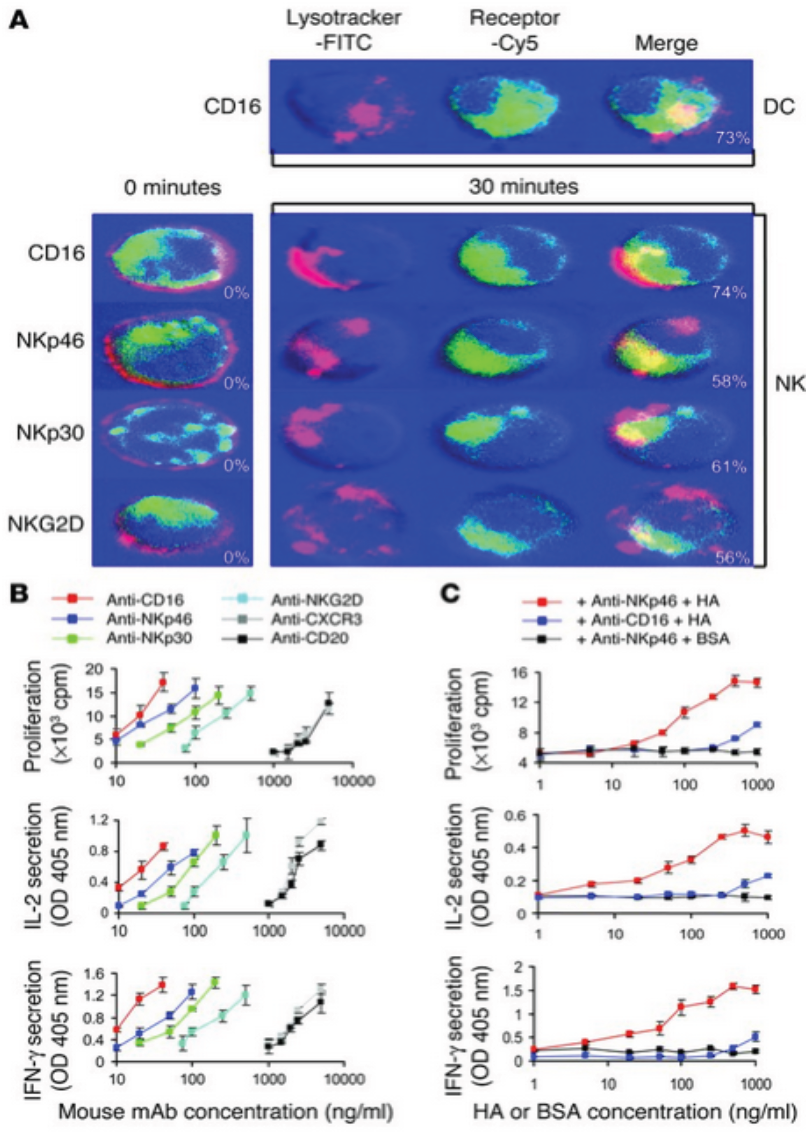
Activating NK receptors internalize into MHC class II peptide loading compartments. **(A)** ANK cells were labeled with LysoTracker and then incubated with anti-CD16, anti-NKp46, anti-NKp30, and anti-NKG2D mAbs and Cy5-labeled F(ab')_2_ goat anti-mouse IgG. NK receptors were visualized by confocal microscopy before (0 minutes) and 30 minutes after induction of internalization at 37_C. CD16 internalization of DCs is shown as a positive control. The lower right corners of merged images indicate the average percentage of cells demonstrating colocalization of NK receptors and endosomal-lysosomal compartments when the cells were visualized at low power. (B) ANK cells were cultured with the indicated mouse mAbs for 24 hours. Subsequently, ANK cells were fixed and introduced to DR4-restricted CD4^+^ T cells specific for C𝓍^40–48^ epitope (50 × 10^3^ cells per well). After 48 hours of incubation, T cell proliferation and cytokine production were measured. (C) Irradiated ANK cells (5 × 10^4^ per well) were washed and incubated with HA or BSA antigens at different concentrations for 30 minutes on ice, then washed and incubated with anti-NKp46 mAb or anti-CD16 mAb for 45 minutes on ice. The cells were further washed and incubated for 30 minutes with goat anti-mouse F(ab')_2_ and washed before addition of 50 × 10^3^ OFR3 clone T cells per well. Cells were cultured for 20 hours and pulsed with [^3^H]thymidine for an additional 16 hours to measure OFR3 T cell proliferation, in addition to IL-2 and IFN-γ production. Data are representative of 3 separate experiments.

To further substantiate that these receptors internalize into MHC class II peptide loading compartments, we performed a classical assay previously used for similar characterization of several receptors on DCs (5, 19). Antibodies can be processed and presented on MHC class II molecules to CD4^+^ specific T cell clones. The presentation of antibody-derived peptides is enhanced if the antibody binds to endocytic receptors that deliver their ligands to intracellular processing compartments (5). Accordingly, we evaluated the ability of ANK cells to present anti-CD16, anti-NKp46, anti-NKp30, and anti-NKG2D mAbs to a C𝓍 -specific HLA-DR4 CD4^+^ restricted T cell clone specific for a peptide epitope derived from mouse IgG1 and IgG2a (5). The presentation of these antibodies was compared with that of an anti–CXCR3 receptor antibody known not to internalize into MHC class II peptide loading compartments (5), and with an anti-CD20 mAb that does not recognize ANK cells. As shown in Figure 6B, antibodies recognizing NK receptors CD16, NKp46, NKp30, and NKG2D were processed and presented to T cells more efficiently than the control antibodies anti-CXCR3 and anti-CD20 (10- to 100- fold depending on the antibody and the concentration used).

Our group has characterized the ability of human NKp46 to recognize HA (6, 20). We therefore questioned whether HA can be internalized following specific recognition by NKp46 receptor and colocalize to MHC class II peptide loading compartments. We exposed NK cells to recombinant HA for a short period of time, washed the nonbound antigen, and cross-linked the NKp46 receptor in order to reduce the pinocytic effect and enhance the amount of HA internalized via the NKp46 pathway. Indeed, this controlled antigen-exposure experiment resulted in overwhelming antigen-specific proliferation and cytokine secretion, evoked at concentrations as low as 20 ng/ml of HA (Figure 6C). The fact that the lower proliferative response was observed only at higher concentrations of HA when anti-CD16 antibody was used suggests that HA-NKp46 complex internalization leads to efficient specific antigen processing and supports the possibility of localization to MHC class II peptide loading endosomal-lysosomal compartments. The lack of enhancement of T cell proliferation when cross-linking of NKp46 was performed without HA protein rules out the possibility that the extensive HA-induced OFR3 T cell proliferation was a result of cytokine secretion that followed NKp46 cross-linking. Taken together, these experiments indicate that NK activating receptors can deliver their ligands to an intracellular compartment where MHC class II loading can occur.

### NK-mediated antigen presentation of immune complexes

Handling of soluble immune complexes by DCs constitutes a unique pathway for antigen acquisition (4). This process is orchestrated via Fc receptors that mediate internalization of antigen-IgG complexes and promote efficient MHC class II antigen presentation. CD16 is widely expressed on the majority of NK cells and is known to facilitate antibody-dependent cytotoxicity. We observed that, like on DCs, cross-linking of CD16 on NK cells leads to its internalization (Figures 5A and 6A). In vitro internalization of CD16 is dependent on the generation of immune complexes (4). We obtained serum samples from 2 healthy donors who received influenza vaccines, and 2 control samples from donors who were not vaccinated. The pooled anti-HA sera were mixed with recombinant HA protein from the same strain of influenza virus, thus forming HA immune complexes. Incubating NK cells with HA immune complexes resulted in a significant specific downregulation of CD16, while HA mixed with pooled control sera, or anti-HA sera mixed with BSA, had no effect (Figure 7A). Furthermore, a significant enhancement in T cell proliferation and cytokine secretion at relatively lower levels of antigen was observed when HA immune complexes were used, and NK cells were at least 5-fold more efficient in antigen presentation in the presence of anti-HA serum (Figure 7B).

**Figure 7.**
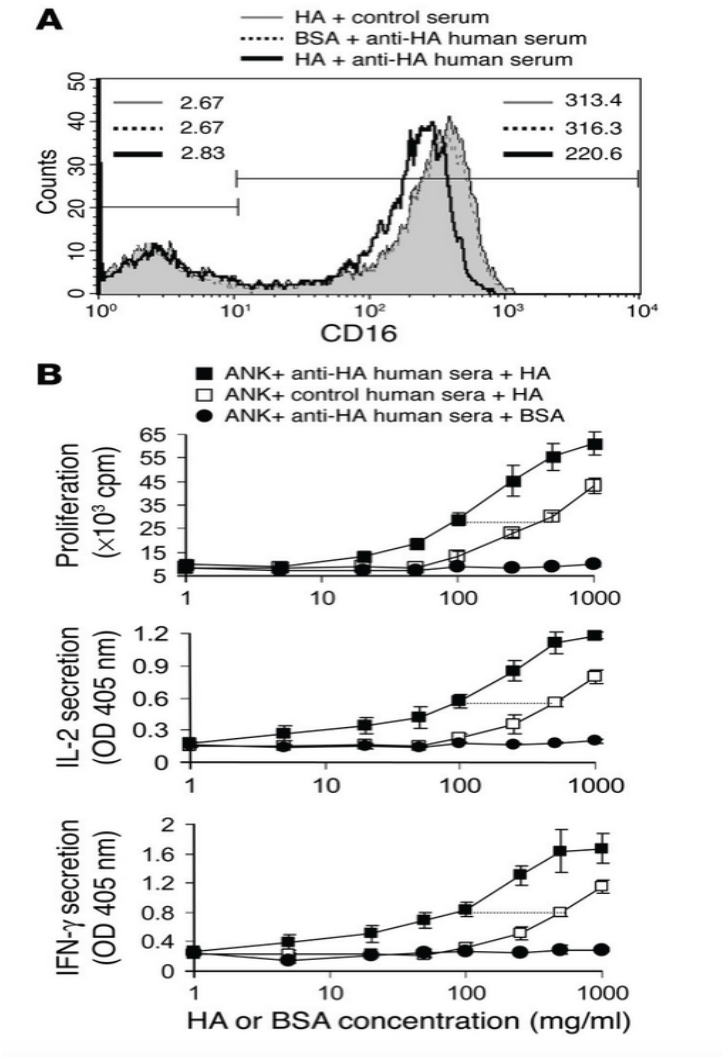
NK-mediated capturing of soluble immune complexes. **(A)** ANK cells (1 × 10^6^) were incubated with recombinant HA or BSA proteins (0.5 μg/ml) and pooled anti-HA or control sera (1 μl per well), for 180 minutes at 37C. The cells were subsequently stained for surface expression of CD16. MFI levels are indicated for CD16^−^ (upper left corner) and CD16^+^ NK cells (upper right corner). (B) The proliferative response of 5 × 10^4^ OFR3 clone T cells to different concentrations of HA or BSA proteins, in the presence of a fixed concentration of pooled anti-HA or control sera (1 μl per well). Irradiated ANK cells (1 × 10^5^) were used as APCs. The cells were cultured for 48 hours and pulsed with [^3^H]thymidine for the last 24 hours. Values are mean ± SD for triplicate samples. Prior to harvesting, 100 μl of supernatant was taken up for ELISA measurement of IL-2 and IFN-γ. Data are representative of 3 separate experiments. In vivo expression of MHC class II and costimulatory molecules on human NK cells. We next sought to characterize the in vivo regulation of MHC class II and costimulatory molecules on human NK cells. We isolated NK cells from inflamed tonsils obtained from 4 donors who underwent elective tonsillectomies. Interestingly, NK cells derived from all samples displayed significant levels of HLA-DR,DP,DQ, CD86, CD70, OX40 ligand, and, less prominently, CD80 (Table 3). These observations indicate that human NK cells acquire APC-like phenotype in vivo in inflamed lymphoid organs without any external manipulation.

**Table 3.** In vivo expression of MHC class II proteins and costimulatory molecules on human NK cells

| Tissue | Sample no. | HLA-DR,<br>DP,DQ (MFI) | CD86<br>(MFI) | CD80<br>(MFI) | CD70<br>(MFI) | OX40 ligand<br>(MFI) |
| --- | --- | --- | --- | --- | --- | --- |
| Chronically inflamed tonsils<sup>A</sup> |  |  |  |  |  |  |
|  | 1 | 108.8 | 71.56 | 11.89 | 66.45 | 68.27 |
|  | 2 | 97.42 | 65.41 | 18.4 | 34.2 | 33.2 |
|  | 3 | 122.3 | 100.2 | 10.2 | 43.4 | 61.42 |
|  | 4 | 134.4 | 87.1 | 11.2 | 69.21 | 48.2 |
| Normal deciduae<sup>B</sup> |  |  |  |  |  |  |
|  | 1 | 13.8 | 4 | 3 | 4 | 4 |
|  | 2 | 11.2 | 3 | 4 | 3 | 4 |
|  | 3 | 11.5 | 4 | 2 | 3 | 4 |
|  | 4 | 12.4 | 4 | 2 | 3 | 3 |
|  | 5 | 14.7 | 4 | 4 | 4 | 3 |
|  | 6 | 9.7 | 4 | 3 | 4 | 3 |
|  | 7 | 11.4 | 3 | 4 | 4 | 4 |
|  | 8 | 10.3 | 3 | 3 | 3 | 4 |
|  | 9 | 8.5 | 4 | 4 | 4 | 4 |
|  | 10 | 11.2 | 4 | 4 | 4 | 4 |
|  | 11 | 13.4 | 7 | 3 | 4 | 3 |
|  | 12 | 11.1 | 4 | 3 | 4 | 3 |
|  | 13 | 14.1 | 3 | 4 | 4 | 4 |
|  | 14 | 11.2 | 4 | 4 | 4 | 4 |
|  | 15 | 9.7 | 4 | 4 | 4 | 4 |
| CMV-infected deciduae<sup>B</sup> |  |  |  |  |  |  |
|  | 1 | 126.1 | 289.3 | 8.08 | 52.29 | 14.9 |
|  | 2 | 134.2 | 122.3 | 12.4 | 48.21 | 33.2 |

<sup>A</sup>Single mononuclear cell suspensions were obtained from chronically inflamed human tonsils and phenotypically analyzed by flow cytometry. CD56<sup>+</sup>CD3<sup>-</sup> NK cells were gated and analyzed for the expression of HLA-DR,DP,DQ, CD80, CD86, CD70, and OX40 ligand. <sup>B</sup>The lymphocyte fraction was isolated from decidual tissue samples from elective terminations of healthy pregnancies (11 samples from first-trimester abortions by uterine scraping, 2 from third-trimester cesarean sections performed with labor, and 2 from third-trimester cesarean sections performed without induction of labor) and from decidual samples from 2 donors suffering from primary CMV infection during pregnancy with evidence of local decidual inflammation and congenital CMV infection of the fetus. The CD3<sup>-</sup> lymphocyte fraction was gated, and decidual NK cells were gated for analysis based on their CD56<sup>+</sup>CD16<sup>-</sup>CD3<sup>-</sup> phenotype and were further analyzed for MHC class II, CD80, CD86, CD70, and OX40 ligand expression. Background MFI values for each staining ranged between 2 and 4.

The delicate balance between immune tolerance and immune activation that prevents immunological rejection of the embryo is maintained via multiple pathways, several of which involve maternal decidual NK (dNK) cells (21). Genomic analysis revealed that these cells can induce decidual immune tolerance via multiple pathways (21). We analyzed the dNK phenotype from 11 decidual samples obtained following first-trimester elective pregnancy terminations, 2 deciduae from third-trimester cesarean sections performed with labor, and 2 deciduae from third-trimester cesarean sections performed without induction of labor. Like UaNK cells, dNK cells expressed low levels of MHC class II and lacked expression of all TCR costimulatory molecules (Table 3). We next searched for clinical cases in which feto-maternal immune tolerance might be drawn out of balance, such as primary maternal CMV infection. Remarkably, dNK cells obtained from CMV-infected mothers expressed high levels of MHC class II and TCR costimulatory molecules (Table 3). Only 2 cases of CMV-infected deciduae were obtained in this study, because primary CMV infection during human pregnancy is rare (8) and its exact detection and diagnosis are quite difficult. Even though no firm conclusion can be drawn, the profound upregulation of these molecules on NK cells obtained from both CMV-infected cases suggests that dNK cells maintain their immunoregulatory potential in normal pregnancy, while they can acquire a proinflammatory phenotype in certain pathologies upon induction of the localized decidual-inflammation cascade.

Impaired TCR costimulatory activity of ANK cells derived from transporter-associated processing antigen 2–deficient patients. NK cells are normally characterized by the ability to lyse cells that lack MHC class I molecules while sparing autologous normal class I–positive cells. Cells derived from patients displaying defective expression of transporter-associated processing antigen 2 (TAP2) are characterized by a strong reduction of MHC class I molecules. This results in the development of severe respiratory manifestations due to recurrent and chronic respiratory infections in these patients. Remarkably, however, in spite of the downregulation of MHC class I molecules, immediate NK cell attack does not occur. NK cells derived from these patients have been extensively studied in order to try to understand the molecular basis underlying this phenomenon. Recent studies suggested that activated NK cells from TAP2-deficient patients developed unique mechanisms to avoid promoting systemic NK-initiated autoimmunity that generally included upregulation of NK-inhibitory receptors and downregulation of activating receptors (22). Several of the activating ligands for TCR costimulatory molecules are major players in propagation and progression of autoimmune disease, providing costimulatory signals for T cells (23). We therefore questioned whether ANK cells derived from TAP2-deficient patients would express lower levels of ligands for TCR costimulatory molecules as part of the general observed NK developmental pattern to avoid NK-initiated autoreactivity. Indeed, ANK cells obtained from all 3 TAP2-deficient patients repeatedly displayed severe impairment in upregulation of CD86, CD80, CD70, and OX40 ligand upon activation (Figure 8). This expression pattern was translated in impaired functional induction of TCR costimulation of autologous total peripheral CD3^+^ T cells in TAP2-deficient patients compared with either their healthy mother (Figure 8) or 12 different unrelated healthy donors (data not shown). Our observations support the notion that an as-yet unknown mechanism promotes the expression of immune-inhibitory receptors on ANK cells from TAP2- deficient patients and attenuates the expression of those that generally promote immune-response progression, such as natural cytotoxicity receptors (NCRs) and T cell–activating molecules.

**Figure 8.**
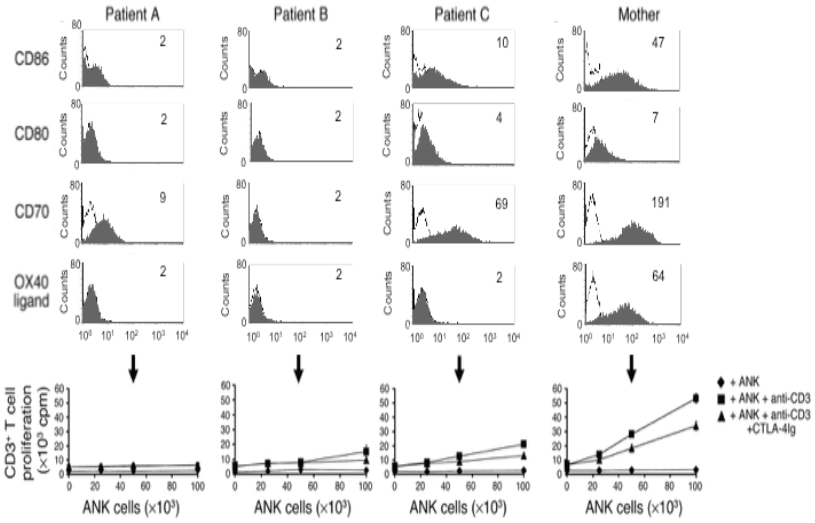
Impaired expression and function of costimulatory molecules on activated NK cells from TAP2-deficient patients. ANK cells were obtained from each of the 3 TAP2-deficient patients and from their healthy mother. Cells were stained by various mAbs as indicated and analyzed by FACS. MFI values for each specific staining (gray histograms) are indicated in the upper right corner. Background MFI values for each staining were between 2 and 4 (white histograms). Bottom panels: Proliferation of purified peripheral blood CD3^+^ T cells from TAP2-deficient patients and their healthy mother, in the presence of irradiated autologous ANK cells with or without suboptimal levels of antibody to CD3 or CTLA-4Ig fusion protein. Values are mean ± SD for triplicate samples. Data are representative of 2 independent experiments.

## Discussion

T cell responses are primed by 3 classes of APCs: DCs, macrophages, and B cells. Each is optimally equipped to present a particular class of antigen to T cells because of distinguished capabilities in antigen capturing (4). DCs are the best professional APCs, as they are highly potent in macropinocytosis and concentrated in peripheral lymphoid and nonlymphoid organs (24). Macrophages are professional scavenger cells that can be induced following pathogen engulfment to present foreign antigens. B cells are uniquely adapted to bind specific soluble molecules through their Ig receptor (2). NK cells, as shown here, are capable of presenting target cell–derived antigens after killing. It is important to emphasize that expression of MHC class II and ligands for T cell costimulatory molecules is not a guarantee for APC capabilities. Eosinophils, for example, cannot process antigens despite significant levels of MHC class II and costimulatory molecules on their surface following activation (25).

Importantly, we demonstrate in vivo that NK cells are able to express MHC class II and TCR costimulatory molecules. Thus NK cells with APC-like phenotype that are present at the right time, in the right place, during the course of the immune response can cross-talk with neighboring cells, including CD4^+^ T cells. Indeed, NK cells can be found in T cell areas of secondary lymphoid organs (spleen, lymph nodes, tonsils) (9, 26) and, as shown here, have the ability to present antigens derived from their target cells after killing (Figure 4). It is clear that DCs are the best APCs; however, in view of our results it is also possible that activated NK cells might “fine-tune” T cell activation and costimulation when direct insult occurs to organs in which they can be found (e.g., direct infection, soluble antigen trafficking via lymph fluid, or tumor metastasis). In addition, the ability of NK cells to localize and become abundant in inflamed nonlymphoid peripheral organs (liver, lungs, intestines, etc.) (7, 9, 26) raises the possibility that these cells can act as important stimulators for effector T cells recruited to these organs at later stages of the immune response (2). The significant levels of MHC class I molecules expressed on NK cells (1), together with ligands for TCR costimulatory molecules, potentially extend the role of NK cells in activating T cells. NK cells are prone to infection by several viruses (1); thus they can potentially assist in evoking pathogen-specific cytotoxic CD8^+^ T cells.

Recently, the induction of a unique regulatory DC subset expressing the NK-specific marker DX5 following anti-CD40 treatment in a mouse autoimmune diabetes model (bitypic DC/NK cells) has been described (27). However, these cells were severely impaired in basic NK cell functions such as perforin expression and cytotoxic capacity. The human NK cells described here seem to differ drastically from mouse bitypic DC/NK cells, as their activation dramatically enhanced both their T cell–activating molecules and their well-regulated lysis (Table 2).

Ligation of CD28 molecule and other costimulatory receptors on T cells promotes their activation and propagation of various types of immune responses, including autoimmunity. NK cells from TAP2-deficient patients are more exposed to activation induction because of the loss of inhibitory signals that results from low systemic MHC class I expression. The functional impairment in the induction of ligands for TCR costimulatory molecules on NK cells derived from these patients upon activation acts in concert with other characterized compensatory mechanisms that aim to keep NK-initiated immune responses under balance (22). On the other hand, the specific induction of these proinflammatory molecules on dNK cells derived from CMV-infected deciduae calls for further elucidation of the involvement of these molecules on NK cells in pathologies in which decidual or graft immune tolerance is compromised.

NK cells are known to kill self cells that are starting to undergo malignant transformation (20). Certain cancer immunotherapy protocols attempt to use NK cells for the clinical treatment of malignancy. For example, infusion of in vitro–activated NK cells back into cancer patients has yielded marked antitumor activity and complete remission in a subset of patients (28). Our observations that NK cells upregulate ligands for TCR costimulatory molecules after killing, localization to inflammatory tissues, or IL-2 treatment might support further attempts to modify such protocols. These molecules can participate in coordinating an effective attack against tumors by engaging multiple immune components, including adaptive T cell responses. There is also growing evidence for the existence of a lymphoid-like microenvironment inside tumors that results in recruitment and local activation of naive CD8^+^ T cells that normally patrol lymphoid tissues (29). It would be interesting to study whether NK cells recruited to these tumor sites might contribute to the creation of this lymphoid platform by acquiring unique antigens from their targets and subsequently directly assisting in MHC-mediated priming of such naive T cells (perhaps specifically against NCR tumor ligands).

Our results directly implicate activating NK receptors in antigen capturing and internalization. T cell and B cell receptor internalization has been implicated in down modulation of TCR responses and in B cell activation, respectively (30). Similarly, ligand-mediated internalization of NK receptors might constitute a mechanism to down regulate NK cytotoxicity during an ongoing immune response and to simultaneously assist in capturing antigens and acquiring APC-like phenotype. Despite the limited characterization of tumor ligands for NCRs, it might be possible to use cultured ANK cells as APCs after they have killed and acquired unique antigens that enable their specific recognition of target tumor cells. Perhaps these cells can present such processed antigens and induce clonal expansion of T cells expressing antigen receptors that can recognize these antigenic epitopes. Such in vitro–expanded specific T cell clones can be potentially utilized for cancer immunotherapy (31).

In summary, we suggest that activated human NK cells are likely to act together with macrophages and B cells as direct key regulators of T cell–mediated immunity, by complementing APC capabilities of DCs. We think that our data unveil an additional layer of complexity in the regulation of human immune response and provide a framework for the pursuit of further study in this new area of NK cell biology.

## Methods

### Cells

Palatine tonsils were obtained from 4 individuals undergoing tonsillectomy for chronic inflammation. Lymphocytes were isolated as previously described (7). Deciduae from elective pregnancy terminations, from induced labors, and from cesarean sections were processed as previously described (8). Primary CMV infection during pregnancy was diagnosed as previously described (8). The TAP2-deficient patients were described previously (22). The institutional review boards of Hadassah Hospital and the Schneider Children’s Medical Center of Israel approved all these studies, and informed consent was obtained according to the World Medical Association Declaration of Helsinki. UaNK and CD4^+^ T cells were purified by negative selection to exclude the presence of contaminants using the NK Cell Isolation Kit II and the CD4^+^ T Cell Isolation Kit II (Miltenyi Biotec Inc.), respectively, according to the manufacturer’s instructions. Purity was routinely checked by FACS analysis and was over 98%. Polyclonal ANK cultures and ImDCs were generated as previously described (10, 20). DC maturation was achieved by addition of LPS (2 ng/ml) 6 hours after incubation with antigen. The HA_306–318_–specific Th clone was generated according to standard protocol (12). Generation of the human CD4^+^ T cell clone that recognizes IgG1- and IgG2a-derived peptide epitope C𝓍^40–48^ has been previously described (5).

### Preparation, culture, and expansion of peptide-specific CD4^+^ T cells

Synthetic peptides HA_306–318_ (PKYVKQNTLKLAT) and HLA-C_124–137_ (LSSWTAADTAAQIT), both known to be HLA-DR4 epitopes (12, 13), were used. HA protein derived from A/Beijing/262/95(H1N1)-like influenza virus was purchased from Protein Sciences Corp. ANK cells were incubated with 150 ng/ml of HA protein or BSA and were subsequently incubated with 4 × 10^6^ autologous CD4^+^ T cells in 24-well plates at a 1:4 ratio. Cells were analyzed on day 10 for detection of peptide-specific CD4^+^ T cells. The protocol for priming of umbilical cord–derived naive CD4^+^ T cells was adapted from a protocol that used cord DCs as APCs (32). Briefly, 4 × 10^6^ purified cord CD4^+^ T cells were incubated with 2 × 10^6^ ANK cells pulsed with HA protein or BSA in 24-well plates. IL-2 (10 U/ml) was added from day 4. Medium was changed every second day after day 7, but no further antigen was added. After 14 days, the cultures were analyzed as indicated in Figure 3E. At day 15, the cultures were restimulated with HA-pulsed ANK cells for 10 days and were subsequently reused as indicated in Figure 3F.

### NK cell membrane preparation

UaNK and ANK cells (100 × 10^6^) were washed thoroughly twice with PBS and then washed twice with homogenization buffer (HEES: 0.255 M sucrose, 2 mM EGTA, 2 mM EDTA, 10 mM HEPES, pH 7.4). Twenty-four milliliters of HEES supplemented with protease inhibitor was added to the cells and was homogenized using a Teflon tissue grinder (Beckman Coulter Inc.). The homogenate was centrifuged, and we next pelleted the plasma membranes by centrifugation of the post nuclear supernatant at 10,500 *g* for 25 minutes. We removed the supernatant, resuspended each pellet in 1 ml of salt wash buffer (0.15 M NaCl, 2 mM MgCl_2_, 20 mM Tris-HCl, pH 7.5), and centrifuged it at 100,000 *g* at 4°C for 30 minutes. The pellet was solubilized in 40 μl of 1% SDS, 10 mM Tris (pH 7.5) solution and boiled at 100°C for 7 minutes. One hundred micrograms of each solubilized membrane fraction was run on 4– 20% Tris-Glycine gel (Invitrogen Corp.). Each resulting separation lane was cut into 20 parts between 250 and 6 kDa. The gel sections were enzymatically digested as previously described (33).

### Mass spectrometry/mass spectrometry analysis of tryptic peptides

The extracted peptides were injected onto a C18 peptide trap precolumn (Michrom Bioresources Inc.) at 100 μl/min. The peptides were chromatographically separated from 5 to 70 minutes with 3–40% acetonitrile/0.1% formic acid at 15 μl/min on a 1 mm × 5 cm, 3-μm C18 PepMap column (LC Packings). The peptides were microsequenced by the mass spectrometer. The mass spectrometry/mass spectrometry spectra were then extracted from the raw data and submitted to the Mascot (34) search engine, available at http://www.matrixscience.com/search_form_select.html, for protein alignment. The Mass Spectrometry protein sequence Data Base (MSDB; http://csc-fserve.hh.med.ic.ac.uk/msdb.html) was used for sequence-matching purposes. The resulting Mascot file was then parsed using scores of 15 for peptides and 20 for proteins.

### Flow cytometry

The following FITC-, PE-, or CyChrome-labeled anti-human mAbs were used: Anti-CD3, anti-CD14, anti-CD20, anti-CD16, anti-CD86, anti-CD80, anti– HLA-DR,DP,DQ, anti-CD70, anti-CD45RO, and FITC- or PE-labeled IgG1 and IgG2a isotype-matched controls were obtained from BD Biosciences — Pharmingen. Anti- CD56 and anti-CD4 were obtained from DAKO Corp. Anti-CXCR3, anti-NKp30, and anti-NKG2D were obtained from R&D Systems Inc. Biotinylated anti–OX40 ligand mAb was obtained from MBL International Corp. Mouse anti–human NKp46 was obtained as previously described (20). PE-labeled DRB1*0401 tetramers loaded with HA or ospA (control) peptide were produced and used as described previously (35).

### RT-PCR analysis

RNA was isolated from UaNK and ANK cells using the RNeasy Mini Kit (QIAGEN Inc.). cDNA was prepared and used for RT-PCR according to standard protocol. B7-H3 primers were 5'-CTGGCATGGGTGTGCATGTG-3', sense, and 5'- ATGGTGACAGAGCCGTGCGC-3', antisense.

### T cell proliferation assays

T cell proliferation assays were performed in 96-well flat-bottom plates in triplicate in a final volume of 250 μl of RPMI plus 10% human serum. In NK costimulation experiments (Figure 2), plates were coated with suboptimal concentrations of anti–human CD3 mAb (T3D, 0.05 μg/ml), washed with RPMI before incubating various numbers of ANK and UaNK cells combined with T cells derived from the same donor. Blocking was applied by Preincubating of NK cells with 25 μg/ml of CTLA-4Ig or CD99Ig fusion proteins before addition of T cells. T cell proliferation was measured by a standard thymidine incorporation assay. Supernatants from T cell proliferation experiments were used for measuring T cell production of human IL-2 and IFN-γ with an ELISA quantikine kit (R&D Systems Inc.). The influenza A/Beijing virus was propagated as previously described (20).

### Internalization assay

A previously described protocol for internalization assay (19) was applied. Internalization was measured by the percentage decrease of mean fluorescence intensity (MFI) compared with that in control samples.

### Confocal microscopy

NK cells were labeled with 500 nM of FITC-LysoTracker (Invitrogen Corp.) for 30 minutes at 37°C, washed, and incubated with mAbs against various receptors for 30 minutes on ice. Samples were next incubated with Cy5-labeled F(ab')_2_ goat anti-mouse IgG for 30 minutes on ice and washed. Cells were observed by a Zeiss Axiovert 100-M inverted microscope (Carl Zeiss Inc.) without incubation or after 30 minutes of incubation at 37°C.

## Manuscript Comment

This manuscript constitutes a corrected version of a paper retracted on April 1^st^, 2015, which was previously peer-reviewed and published in the Journal of Clinical Investigation by our group (Novel APC-like properties of human NK cells directly regulate T cell activation- **J Clin Invest. 2004;114(11):1612–1623. doi:10.1172/JCI22787**). This version of the manuscript, now published on bioRxiv, includes the following corrections that have not been peer-reviewed, but rather validated and/or newly generated by our group:

1. ANK and UaNK samples were mislabeled in the gel presented in Figure 1 in the original manuscript, which is now corrected, and we add red dotted line to highlight cropping. All other gels accurately reflect the experiments described as originally presented.
2. We realized that the FACS plots in Figure 8 were inadvertently generated from the wrong FACS experiment data folder (which involved many of the same surface markers – OX40ligand, CD70, CD86, CD80 - used to generate Figures 3 and 6 in Hanna et al. J Immunology. 2004 Dec 1;173(11):6547-63). This mistake happened by our team, while the first author had limited access to the data while being abroad and away from the lab writing and revising all three manuscripts simultaneously. To resolve this, we have validated and repeated this experiment and now provide a corrected Figure 8. (We note that all Ethical and Helsinki Approvals and Board permits are still maintained and valid to conduct these experiments).
3. Due to the same issue, 2 negative control FACS plots in Figure 3D were inadvertently misrepresented from the wrong FACS files during revision of this paper (+ HA (middle of bottom row), UaNK + HA (in right of upper row)). We repeated the experiment presented in Figure 3D, and a corrected Figure is now included where we chose to provide 2 validated and representative control FACS panels (rather than replace the entire panel, in order to keep changes to a minimum).

We apologize for this inconvenience and the mistakes in the original JCI manuscript (J Clin Invest. 2004;114(11):1612–1623. doi:10.1172/JCI22787), which we revalidate and correct herein. All conclusions reported in the original manuscript remain corroborated and valid following this correction.

## Acknowledgements

This research project was funded by research grants from the Israel Cancer Research Foundation, the Israel Science Foundation, the European Commission, the Hadassah Women’s Health Center, and the Fritz Thyssen Foundation. We thank all of the authors of the original published version of this manuscript (Hanna et al. J Clin Invest. 2004;114(11):1612–1623. doi:10.1172/JCI22787).

